# AMP-activated protein kinase protects against anoxia in *Drosophila melanogaster*

**DOI:** 10.1101/162719

**Authors:** Justin J. Evans, Chengfeng Xiao, R. Meldrum Robertson

**Author notes:** Corresponding author: RMR.

## Abstract

During anoxia, proper energy maintenance is essential in order to maintain neural operation. Starvation activates AMP-activated protein kinase (AMPK), an evolutionarily conserved indicator of cellular energy status, in a cascade which modulates ATP production and consumption. We investigated the role of energetic status on anoxia tolerance in *Drosophila* and discovered that starvation or AMPK activation increases the speed of locomotor recovery from an anoxic coma. Using temporal and spatial genetic targeting we found that AMPK in the fat body contributes to starvation-induced fast locomotor recovery, whereas, under fed conditions, disrupting AMPK in oenocytes prolongs recovery. By evaluating spreading depolarization in the fly brain during anoxia we show that AMPK activation reduces the severity of ionic disruption and prolongs recovery of electrical activity. Further genetic targeting indicates that glial, but not neuronal, AMPK affects locomotor recovery. Together, these findings support a model in which AMPK is neuroprotective in *Drosophila*.

## Introduction

Optimal neural function requires an uninterrupted supply of energy. Prolonged disruption of this supply can induce neuronal damage, whereas short periods can be tolerated and some organisms are more tolerant by virtue of adaptive mechanisms of metabolic suppression (Staples and Buck, 2009). *Drosophila* survive anoxic conditions, which induce severe energetic depletion, by entering a reversible coma (Lighton and Schilman, 2007; Benasayag-Meszaros et al., 2015). The coma is generated by anoxic depolarization in the brain (Armstrong et al., 2011) which is a form of spreading depolarization (SD), a loss of CNS ion gradients common to insects and mammals (Spong et al., 2016a) and associated with several important brain pathologies (Dreier and Reiffurth, 2015). During this coma, survival depends on the down-regulation of energy turnover and up-regulation of ATP-producing pathways (Hochachka et al., 1996). An enzyme known to play a role in coordinating energy allocation is the metabolic sensing protein, adenosine monophosphate-activated protein kinase (AMPK). In insect preparations AMPK activation mimics hypoxia and starvation by reducing neuronal excitability (Money et al., 2014) and exacerbating ouabain-induced SD (Rodgers-Garlick et al., 2011). Activation of AMPK contributes to mechanisms for coping with hypoxia by metabolic suppression in non-mammalian vertebrates, such as fish (Jibb and Richards, 2008; Zhu et al., 2013). AMPK has diverse roles in different mammalian tissues (Mantovani and Roy, 2011) but, in spite of much research on the role of AMPK in neural damage after oxygen limitation associated with stroke (Manwani and McCullough, 2013) there is little knowledge of its potential role in modulating induction and recovery from SD.

AMPK is a heterotrimeric protein consisting of a catalytic alpha (α) subunit, regulatory gamma (γ), and scaffolding beta subunit (β). Allosteric activation by AMP leads to the phosphorylation of the α subunit on a threonine residue (Thr-172) (Hawley et al., 1995). A rise in AMP, via the adenylation kinase reaction, occurs when ATP is depleted. Since AMP:ATP varies as a square of ADP:ATP, the AMPK cascade is a sensitive indicator of cellular energy change (Hardie et al., 1998). Once activated, AMPK inactivates the ATP-consuming anabolic pathways: fatty acid synthesis via phosphorylation of acetyl-CoA carboxylate and sterol synthesis via phosphorylation of 3-hydroxy-3-methylglutaryl-CoA reductase (Hardie et al., 1999). As a result, depression of acetyl-CoA carboxylate activates catabolic ATP-production via a reduction in malonyl-CoA concentration. Low concentration of malonyl-CoA reduces inhibition of carnitine palmitoyltransferase I (CPT-I), leading to an influx of fatty acid substrate (Guzman and Blazquez, 2004). Surplus fatty acid is converted into ketone bodies to be used as a source of alternative cellular energy during dietary restriction or prolonged starvation (Murray et al., 2016).

Mechanisms of metabolic regulation during dietary restriction are highly conserved. In mammals, dietary restriction leads to the accumulation of lipids in the liver, which are later oxidized to become ketone bodies. While ketone bodies are mainly synthesized and supplied by the liver, studies suggest glia exhibits hepatic-like ketogenic machinery (Blazquez et al., 1998). Dietary restriction, or 'ketogenic diets,' can protect the brain from oxidative stress in mammals as well as in *Drosophila* (Suzuki et al., 2001; Vigne et al., 2009; Gibson et al., 2012). In *Drosophila,* the oenocytes and fat body have metabolically similar liver-like functions (Gutierrez et al., 2007). Similarly, dietary restriction increases lipid accumulation in oenocytes and is associated with heightened levels of ketone bodies in the brain (Johnson et al., 2010; Schulz et al., 2015). While AMPK is activated in periods of dietary restriction (Slack et al., 2012), the role of AMPK in these tissues during anoxic stress is still unknown.

Here we show that AMPK modulates recovery from anoxic comas in *Drosophila*. This process involves the return of ion homeostasis in the brain followed by recovery of neural circuit function upon return to normoxia. We found that pharmacological AMPK activation mimics the effects of dietary restriction in regulating anoxic recovery and demonstrated using genetic manipulations that AMPK function in glia, rather than neurons, is important for recovery of locomotor circuits from SD. In addition, AMPK activity in tissues that drive metabolism (oenocytes, fat body, and glia) is central to recovery from anoxia. Our results are consistent with models that suggest AMPK mediates the supply of alternative neural energy and support the conclusion that the AMPK cascade is neuroprotective in *Drosophila*.

## Material & Methods

### Drosophila culture

All control lines were maintained on standard agar medium (SAM): 0.01 % molasses, 8.20 % cornmeal, 3.40 % killed yeast, 0.94 % agar, 0.18 % benzoic acid, 0.66 % propionic acid and 86.61 % water at room temperature (25 ± 0.25 °C) and 60-70 % relative humidity. Emerging flies were collected over a 48 hour period using nitrogen (N_2_) anesthesia and transferred to new food vials. All vials were maintained at equal densities (approx. 20 flies). Experiments were performed on male flies aged 5-9 days after eclosion. Flies were given a minimum of three days without N_2_ exposure before experimentation to mitigate acute tolerance to anoxia.

### Fly strains

*w*^*1118*^ flies were used as control. Tissue specific manipulation of AMPK was achieved using the Gal4/UAS system. The UAS lines used to upregulate, UAS-Cherry-AMPKα (#51871), and downregulate, UAS-AMPKα-RNAi (#25931), AMPK have been previously described (Swick et al., 2013; Li et al., 2016).

GAL4 enhancer-trap strains with broad tissue expression included the pan-neuronal (*Elav* #8765) and pan-glial (*Repo* #7415) lines. Specific glial expression lines included surface glia (subperineural: *NP2276*) and neuropil glia (astrocyte: *Alrm II, Alrm III, NP1243*; ensheathing: *mz0709, NP6520*) (see (Awasaki et al., 2008) for a detailed review). Transgene controls were backcrossed to the *w*^*1118*^ background.

Expression patterns of mifepristone (RU486) steroid-activated Gal4 lines used are: P[Switch2]GSG3448 in fat body, oenocytes, and tracheal cells; P[Switch2]GSG10751 in fat body; P[Switch2]GSGB9-1 in oenocytes. All GSG lines were generated and described by Nicholson *et al.* (2008), with expression patterns shown in supplemental data (http://flystocks.bio.indiana.edu/Browse/gal4/gal4_switch.php).

All lines were obtained from the Bloomington Stock center (Bloomington, IN).

### Treatments

#### Starvation

In preliminary experiments, flies exhibited optimal survival and anoxic tolerance after 24 hrs of starvation; *w*^*1118*^ flies were tested at 12, 24, 36 hours and during a survival assay (data not shown). Under starvation conditions, flies were transferred 24 hours before testing into vials containing water-soaked filter paper. The vials were rehydrated 12 hours into starvation to mitigate desiccation effects.

#### (B) Pharmacology

Metformin (1-1 dimethylbiguanide hydrochloride; Sigma-Aldrich) was added directly into heated standard agar medium, to make a 100 mM concentration, and allowed to settle for 24 hours before testing. AICAR (5-aminoimidazole-4-carboxamide-1-beta-D-ribose-furanoside; TOCRIS) was dissolved into 1% dimethylsulfoxide (DMSO) to make a 100 mM aqueous stock. Approximately 250 μl aliquots, from the stock, were layered on the standard agar medium and allowed to absorb for 24 hours before testing (Vigne et al., 2009). Controls were layered with 250 μl of 1% DMSO. Flies were fed AICAR or metformin for four days prior to testing.

#### (C) Gene-Switch System

Gene-Switch flies were fed RU486 (mifepristone; Sigma-Aldrich) based on optimal time and dose-dependent transcriptional activation reported (Osterwalder et al., 2001). RU486 was mixed into heated standard agar medium (10 μg/mL) and allowed to settle for 24 hours before testing. Gene-Switch flies under non-starvation conditions were placed on either standard agar medium or standard agar medium with RU486 for two or four days prior to testing. Gene-Switch flies under starvation conditions were fed standard agar medium for at least four days and then transferred 24 hours before testing into vials containing either non-nutritive agar or non-nutritive agar with RU486 (10 μg/mL).

### Locomotor Assay

A high-throughput locomotor assay (Xiao and Robertson, 2015) was used to evaluate changes in activity level before and after anoxia. Flies were loaded into circular chambers (1.27 cm diameter x 0.3 cm height) (Fig. 1A); one fly per chamber. Video recording (15 fps) was used to capture two-dimensional locomotion as the chamber restricts vertical movement (Video 1S). Fly tracking software (Xiao and Robertson, 2015), was used to calculate each fly's location every 0.2 s throughout the recording period. The positions (x, y) of each fly was exported to Excel spreadsheets to evaluate different parameters.

**Figure 1.**
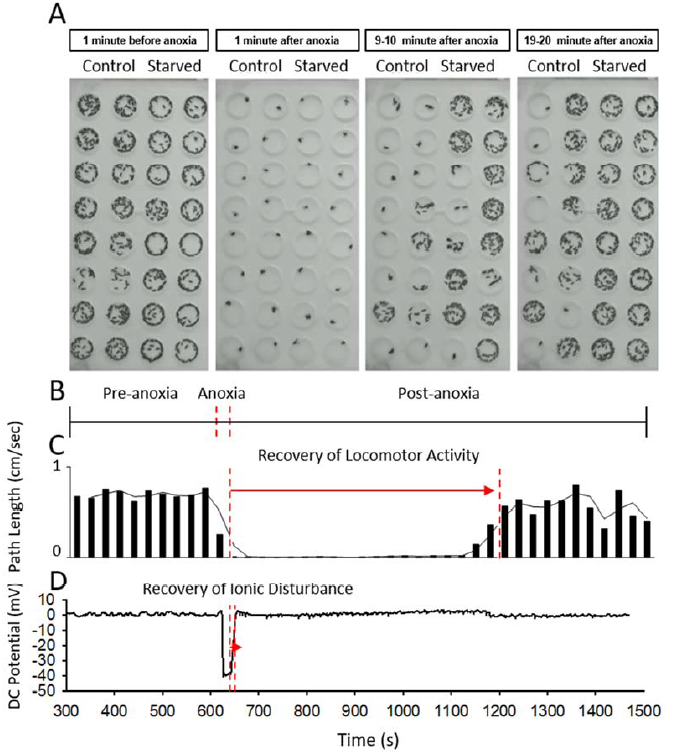
Locomotor recovery from anoxic coma. **(A)** Behavioral assay. Superimposed images (300) from 1 minute of video recording at 4 different time points before, during and after an anoxic coma. Each chamber contained a single fly. Note that starved flies recovered locomotion faster after an anoxic coma. See supplemental video, **Video 1S**. **(B)** The recording period includes baseline activity (10 min), anoxia (30 s of N_2_), and post-anoxic activity (60 min). **(C)** Locomotor recovery of an individual fly indicated by average path length (cm) every 30 seconds (columns) and a moving average of path length (line). Timing of anoxia aligned with B. **(D)** Extracellular recording of DC potential (mV) from the brain. Abrupt negative DC shift indicates spreading depolarization. Note that recovery of ion gradients occurs rapidly in contrast to recovery of locomotion. Timing of anoxia aligned with B and C. 250

The total recording included periods of baseline activity (10 min), anoxia (30 s of N_2_), and post-anoxic activity (60 min) (Fig. 1B). During the experiment airflow was maintained at 2 L/min to avoid hypoxic conditions, and a N_2_ (8 L/min) exposure was used to induce an anoxic coma. To allow animals to recover from handling, the first five minutes of each baseline recording was disregarded (Liu *et al.* 2007). Further, as placement in the chambers can injure the flies, individuals that failed to recover 10% of regular baseline activity (250 cm / 5 min) following anoxia were excluded from further analysis.

### Locomotor Activity Parameters

#### (A) Pre-anoxia

Pre-anoxia parameters were used to evaluate changes in locomotor activity during the baseline period. Path Length (PL) was used to measure the total distance travelled (cm) during the 5 minute pre-anoxia period. Time at Rest (TAR) denotes a time period, 2 seconds or longer, during which the individual did not meet the defined 'activity' threshold (0.15 cm per second). This 'inactive' time period was recorded as a percentage of the pre-anoxia time period. The rationale was that both PL and TAR could be used to evaluate the hyperactivity commonly associated with starvation (Lee and Park, 2004; Johnson et al., 2010).

#### (B) Anoxia & Post-anoxia

Parameters during and after anoxia were used to characterize changes in susceptibility and tolerance to anoxic conditions. The parameters which evaluated susceptibility to anoxia (30 s of N_2_) include Time to Succumb (TS) (Fig. 2Aii). TS was measured as the time (s) from N_2_ onset to the point when activity fell below the recovery threshold (< 0.30 cm per second). Recovery from anoxia, denoted Time to Recovery (TR), was measured as the time taken for locomotor activity to surpass a defined threshold (0.3 cm per second) at least 10 times within 60 consecutive seconds (Xiao and Robertson, 2016). The earliest time to reach the criterion is set as TR (Fig. 2A). As normal TR occurs within 30 minutes, individuals that failed to meet the TR criterion within the 60 minute post-anoxia period were excluded in order to make comparisons between 'recovered' individuals.

**Figure 2.**
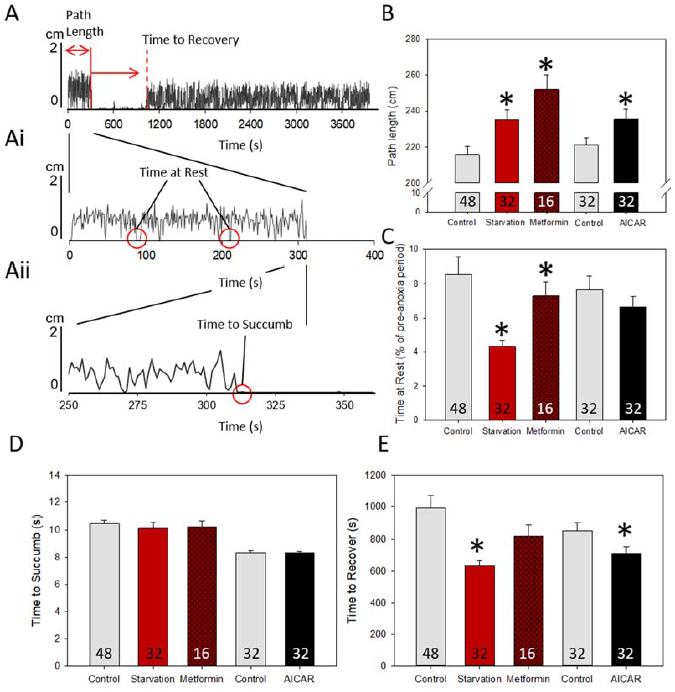
Starvation and AMPK activation increase locomotor activity and speed recovery from an anoxic coma. **(A) Ai** Locomotor activity parameters. Path Length (PL) is the distance (cm) traveled during the pre-anoxia period. Time at Rest (TAR) is the percentage of time during the pre-anoxia period that the individual does not meet the activity threshold (< 0.15 cm/s) for periods of at least 2 s. Time to Recovery (TR) (s) indicated by the dashed line. **Aii** During anoxia (30 s of N_2_), the Time to Succumb (TS) is the time (s) from nitrogen on to a period of activity below the recovery threshold (< 0.30 cm/s). **(B, C)** 24 hrs of starvation significantly increased PL and decreased TAR. Pharmacological activation of AMPK by feeding 100 mM metformin or 100 mM AICAR for four days mimicked the effects of starvation. p < 0.05 (*) by *t*-test or Mann-Whitney Rank Sum. **(D)** Treatments did not affect TS. **(E)** Starvation and AICAR significantly reduced TR. p < 0.05 (*) by Mann-Whitney Rank Sum. Data are plotted as mean ± SE. Sample sizes are indicated in the bars. Starvation and metformin controls are combined for analysis; significance comparisons were made between treatments and their respective controls.

### Spreading Depolarization: DC potential shifts

Flies were placed head first into a disposable pipette tip, without the use of N_2_, and fixed in place using a small drop of wax. Using microscissors, a small window (0.06 × 0.02 mm) was made in the cuticle at the back of the head behind the ocelli (Rodriguez and Robertson, 2012). A chlorided silver wire was placed into the mid-thorax to ground the preparation. Next, a microelectrode pulled from a 1 mm diameter filamented glass capillary tube, backfilled with 1mM KAC, was inserted at an approximate 45º angle through the window into the hemolymph. Further insertion of the microelectrode, from the hemolymph to the brain, was characterized by a negative shift in the resting DC potential.

SD in the fly brain was recorded by monitoring DC potential shifts using a Model 2000 pH/ion amplifier (World Precision Instruments Inc) using the electrophysiology software Clampex for later analysis using Clampfit (Molecular Devices, version 10.3). The recording period included periods of baseline activity (10 minutes), anoxia (30 seconds of N_2_), and post-anoxia (20 minutes). Individuals that exhibited spontaneous, large amplitude DC shifts, which can result from damage, during the baseline recording period were discarded. Direct airflow was maintained throughout the experiment and N_2_ was used to induce an anoxic coma.

### Spreading Depolarization Parameters

The baseline of each recording was adjusted to zero just before the start of anoxia (10 minutes) and after the major ionic disruption in order to normalize DC potential drift during the recording. The time to surge (s) was measured from the time nitrogen was turned on to the point at half-max amplitude of the abrupt negative shift in DC potential. The time to recover (s) was measured from the time nitrogen was turned off to the point of half-max amplitude on the positive shift in DC potential. The peak amplitude (mV) during the anoxia was used to quantify the magnitude of spreading depolarization.

### Measurement of Mass

Adult flies were placed in a freezer (~30 min) before weighing to ensure complete immobilization, after which they were individually weighed to 1 μg using a Cahn-microbalance. Controls were weighed at the same time of day to account for potential water loss. Pupae were weighed under similar conditions. Each pupa was dissected under a light-microscope to verify sex, and genotype via eye color.

### Statistical Analyses

Each locomotor activity test included all controls; for example both Gal4/+ and UAS/+ flies for genetic manipulations or drug-specific controls for flies fed AMPK pharmacology. As replicate data sets exhibited similar trends we compiled the data for statistical analyses.

Statistical analyses were performed using Sigma Plot 12.1 (Systat Software Inc.). Parametric tests performed were a t-test and one-way ANOVA when the distribution of residuals passed normality (Shapiro-Wilk). Non-parametric tests, when normality failed, included a Mann-Whitney Rank Sum Test and ANOVA on Ranks (Kruskal-Wallis) with Dunn's multiple comparisons. Significance levels were set at α < 0.05 and 0.10.

## Results

### AMPK activation mimics starvation phenotype before and after anoxia

To evaluate phenotypic changes in locomotor behavior before, during, and after exposure to anoxia we used a high-throughput locomotor assay and open-source software to track each fly's location (Fig. 1; see also Video 1S) (Xiao and Robertson, 2015). Flies placed within the restricted locomotor chamber engaged in near constant walking (Fig. 2) [Time at Rest (TAR): 8.7 ± 2.1 %; Path Length (PL): 215.3 ± 6.5 cm]. When deprived of food for 24 hrs the flies exhibited a distinct 'starvation phenotype'. Starvation significantly decreased TAR 4.3 ± 0.36 % and increased PL 235.2 ± 5.32 cm (*Mann*-*Whitney, U* (31, 32) = 1184.5, p = 0.008; *Mann*-*Whitney, U* (31, 32) = 827, p = 0.024). When fed the AMPK activating compounds metformin and AICAR, w1118 flies exhibited an increase in PL (Metformin: *t*-test, *t* = −3.355, *df* = 30, p =0.001; AICAR: *t*-test, *t* = −2.088, *df* = 62, p =0.021), however only metformin decreased TAR (Metformin: *Mann*-*Whitney, U* (16, 16) = 62.0, p = 0.013; AICAR: *Mann*-*Whitney, U* (32, 32) = 460.5, p = 0.493). This suggests that, under normoxic conditions, pharmacological activation of AMPK has a similar effect on locomotor activity as starvation.

When exposed to brief anoxia (30 s of N_2_), w^1118^ flies undergo stereotyped movements: rapid paralysis, falling over, and inactivity. We found that the time for flies to exhibit inactivity (Time to Succumb: TS) after exposure to anoxia was not significantly different after a 24 hr period of starvation, or when fed Metformin or AICAR (Fig. 2D) (Starvation: *Mann*-*Whitney, U* (31, 32) = 463.0, p = 0.649; Metformin: *Mann*-*Whitney, U* (16, 16) = 113.0, p = 0.575; AICAR: *Mann*-*Whitney, U* (32, 32) = 1072.0, p = 0.631). Upon return to normoxia, w^1118^ flies starved for 24 hrs or fed AICAR had a significantly shorter time to recover locomotion (Time to Recover: TR) (Starvation: *Mann*-*Whitney, U* (25, 32) = 253.0, p = 0.018; AICAR: *Mann*-*Whitney, U* (32, 32) = 360.0, p = 0.042). While flies fed metformin showed a similar trend, treatment did not significantly affect TR (*Mann*-*Whitney, U* (16, 16) = 89.0, p = 0.147). To determine if this was due to neural mechanisms, we next investigated the effect of starvation and pharmacological activation of AMPK on ionic disruption in the brain during anoxia.

To determine if AMPK could modify CNS tolerance to anoxia we induced a 30 s anoxic coma and monitored spreading depolarization (SD) in the brain. SD refers to the loss of ion gradients induced by a failure of ion homeostasis. The rise in extracellular K^+^ and other ion disturbances induce a large negative shift in extracellular DC potential causing the invertebrate CNS to abruptly shut down in response to acute stress (Rodriguez and Robertson, 2012; Spong et al., 2016b). Anoxia induces SD in the fly brain, pre-anoxia positivity (1 s) and a large negative DC shift (surge) which is sustained until N_2_ is turned off (Fig. 3A), that is similar to anoxic depolarization in the mammalian brain (Hansen, 1985; de Crespigny et al., 1999; Lindquist and Shuttleworth, 2017). We found that the time to CNS shutdown, defined as the time to half-max amplitude of the surge (s), was significantly longer when flies were starved 24 hours or fed metformin (Fig. 3B) (Starvation: *t*-test, *t* = −1.981, *df* = 19, p =0.03); Metformin: *t*-test, *t* = −2.298, *df* = 20, p =0.016). However, flies fed AICAR, which acts as a more direct activator of AMPK, did not exhibit a longer time to surge (*Mann*-*Whitney, U* (6, 10) = 25.0, p = 0.625). We next asked if starvation or AMPK pharmacology would affect the magnitude of SD in the fly brain.

**Figure 3.**
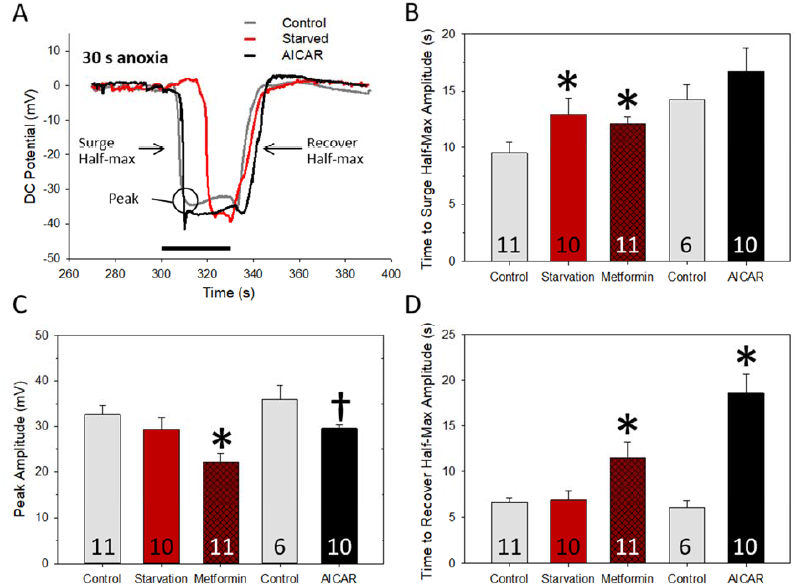
Starvation and AMPK activation have similar effects on anoxia-induced SD in the brain. **(A)** Representative extracellular recording of DC field potential showing abrupt negative shifts induced by anoxia in flies: control (grey), after 24 hrs starved (red), or fed 100 mM AICAR (black). The Peak (mV) of the DC shift was used to assess the magnitude of disruption. Time to Surge Half-Max Amplitude (s) and Time to Recover Half-Max Amplitude (s) were measured to assess the rate of failure and clearance mechanisms. **(B)** Time to Surge Half-Max Amplitude was longer in flies starved 24 hrs or fed 100 mM metformin. p < 0.05 (*) by *t*-test or Mann-Whitney Rank Sum. **(C, D)** Metformin or AICAR reduced peak amplitudes and increased Time to Recover Half-Max Amplitude. p < 0.05 (*), 0.10 (ƚ) by *t*-test or Mann-Whitney Rank Sum. Data are plotted as mean ± SE. Sample sizes are indicated in the bars. Starvation and metformin controls are combined for analysis; significance statistical comparisons were made between treatments and their respective controls.

We predicted that flies with faster locomotor recovery would have smaller ionic disturbances during anoxia, enabling them to re-establish ionic gradients across cell membranes faster. What we found was that during anoxia, the peak amplitude was significantly smaller in flies fed Metformin or AICAR, but not after a 24 hr starvation period (Fig. 3C) (Metformin: *Mann*-*Whitney, U* (11, 11) = 22.0, p = 0.013; AICAR: *Mann*-*Whitney, U* (6, 10) = 14.0, p = 0.093; Starvation: *t*-test, *t* = 0.975, *df* = 19, p =0.17). However, following anoxia, flies emerging from an anoxic coma, defined as the time to recover half-max amplitude (s), took significantly longer when fed Metformin or AICAR (Fig. 3D) (Metformin: *Mann*-*Whitney, U* (11, 11) = 16.0, p = 0.004; AICAR: *Mann*-*Whitney, U* (6, 9) = 0.0, p = 0.002), while flies starved 24 hrs did not exhibit a difference (*Mann*-*Whitney, U* (10, 11) = 51.0, p = 0.805).

Thus, contrary to what we expected, fast locomotor recovery did not correlate directly with smaller ionic disturbance and flies with a smaller peak amplitude took longer to recover CNS ion gradients after anoxia.

Although we have shown that both starvation and the AMPK activators Metformin and AICAR were able to modulate SD dynamics in the brain during anoxia, it was unclear whether the AMPK modulation of the speed of locomotor recovery after anoxia was mediated neurally. We tested this possibility using the *Drosophila* Gal4/UAS system to up- or downregulate the AMPK catalytic α subunit activity in a tissue-specific manner.

### AMPKα subunit required for pupation

We suspected that AMPK would be required during development as *Drosophila* larvae forage in a hypoxic medium (Callier et al., 2015). Complete loss of AMPK*α,* as seen in *ampkα* mutants, is lethal in late second/third instar stages (Lee et al., 2007; Swick et al., 2013). Thus, we chose to restrict RNAi manipulation of AMPK*α* to neurons or glia. We were intrigued to find that a panneuronal up- or down-regulation of AMPK*α* did not affect developmental lethality, locomotor behavior, or recovery from anoxia (Fig. 4, Table 1S). In contrast, we found flies with panglial upregulation of AMPKα had a shorter anoxic recovery time (Figure 4B) (Student-Newman-Keuls Method, Q = 3.545, 3.754, p < 0.05), however we were unable to test the effect of pan-glial AMPKα downregulation as these flies showed a male-specific lethality during pupation. As *dAMPKα* mutants exhibit low triglyceride levels, small fat body cells, and decreased larval size (Bland et al., 2010) we evaluated potential weight differences during pupation. We found that there was no weight difference in glial AMPK-RNAi male or female flies during pupation, consistent with a weight-independent lethality during pupation (Fig. 4F) (male pupae: ANOVA, p > 0.05; female pupae: Dunn's Method, Q = 1.926, 1.399, p > 0.05). This suggests that AMPK*α* may be required in glial cells to regulate pupal metamorphosis. In order to circumvent this lethality we used glial subtypes to drive AMPKα-RNAi.

**Figure 4.**
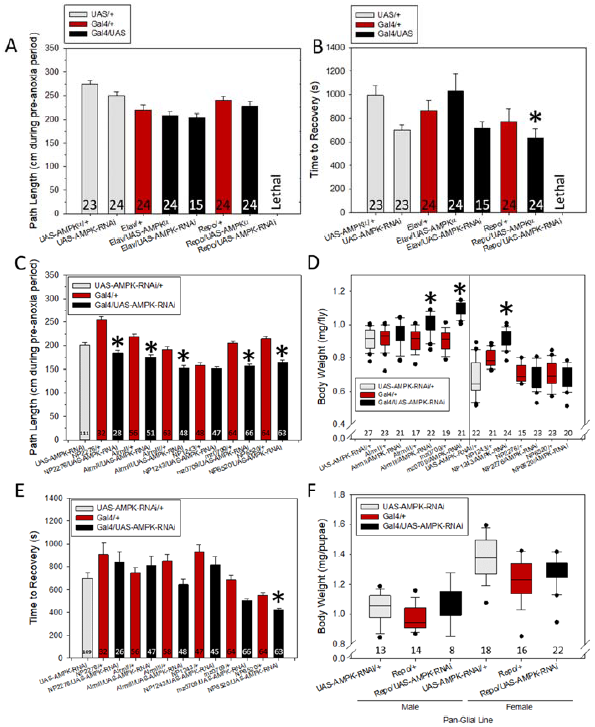
Effects of manipulation of AMPKα subunit expression in neurons and glia. **(A)** Prolonged (6-8 day adults) upregulation or downregulation of the AMPKα subunit in pan-neuronal (*Elav*) or pan-glial (*Repo*) tissue had no effect on path length (PL). Pan-glial downregulation of the AMPKα subunit was lethal in larval stages. **(B)** Pan-glial upregulation of AMPKα significantly reduced TR. p < 0.05 (*) by Dunn’s Method. **(C, D)** Glial-specific downregulation of the AMPKα subunit was investigated in surface (subperineural: *NP2276*) and neuropil (astrocyte: *Alrm II, Alrm III, NP1243*; ensheathing: *mz0709, NP6520*) glia. Downregulation of the AMPKα subunit significantly reduced PL in all glial-specific lines except *NP1243*. Downregulation in the astrocyte lines (*Alrm III, NP1243*) and ensheathing glial line (*mz0709*) significantly increased body weight. The black line in the middle indicates fly line weight was measured at separate times (controls specific to each group). p < 0.05 (*) by Dunn’s Method or Holm-Sidak comparisons. **(E)** The ensheathing glial line *NP6520* reduced TR after anoxia. p < 0.05 (*) by Dunn’s Method. **(F)** Pan-glial AMPKα downregulation had no effect on male or female body weight. Data are plotted as median ± upper and lower quartiles or mean ± SE. Sample sizes are indicated in bars or below boxplots.

### Inhibition of glial AMPKα subunit induces severe metabolic defects

Despite pan-glial AMPKα-RNAi lethality, males with AMPKα-RNAi in surface glia (subperineural: *NP2276*) or neuropil glia (astrocyte: *Alrm II, Alrm III, NP1243*; ensheathing: *mz0709, NP6520*) survived to adulthood. We found that RNAi of AMPKα in the neuropil glial *Alrm III, NP1243, mz0709* increased body weight (Fig. 4D; *Alrm III*: Holm-Sidak ANOVA, t (2) = 4.932, 4.639, p < 0.001; *NP1243*: Dunn's Method H (2) = 44.033, p < 0.05; *mz0709*: Holm-Sidak ANOVA, t (2) = 11.310, 10.852, p < 0.001). When we evaluated locomotor behavior, we also found that nearly all glial subytpes exhibited reduced locomotor activity (Fig. 4C; *Alrm II*: Holm-Sidak ANOVA, t (2) = 5.167, 3.544, p < 0.001; *Alrm III*: t (2) = 6.519, 4.743, p < 0.001; *mz0709*: Dunn's Method, Q = 6.479, 6.630, p < 0.05; *NP2276*: Holm-Sidak ANOVA, t (2) = 6.878, 2.079, p < 0.05; *NP6520*: t (2) = 6.846, 2.012, p < 0.05). While it is unclear whether reduced locomotor activity in the AMPKα-RNAi glial lines was due to increased body weight, it is clear that a loss of AMPKα in glia severely disrupts regular metabolic function.

As a higher rate of metabolism during anoxia could more rapidly deplete ATP required to maintain homeostasis (Galli and Richards, 2014), we were not surprised that a low metabolic rate phenotype - high body weight and low activity level -was associated with a significantly faster anoxic TR in the ensheathing glial line *NP6520*/UAS-AMPKα-RNAi (Fig. 4E; Dunn's Method, Q = 5.413, 4.189, p < 0.05). While *NP6520*/UAS-AMPKα-RNAi did exhibit a significantly faster TS than its controls, a phenotype not observed in the other glial subtypes, variations in TS were not associated with TR (Table 1S).

### Gene-Switch line starvation phenotype

We utilized temporal and spatial downregulation of AMPKα in order to identify AMPK-dependent tissues within the fly body during starvation. We chose to target AMPKα-RNAi in the fat body, oenocytes, and tracheal cells which become transcriptionally inactivated when flies are fed RU486 (mifepristone; Sigma-Aldrich) using Gene-Switch system lines, and scored starvation-induced lethality over an 84 hour period (Osterwalder et al., 2001). Consistent with research that shows pan-neuronal upregulation of AMPK shortens survival during starvation (Ulgherait et al., 2014), we found that AMPKα downregulation in the GSG3448 (fat body, oenocytes, and tracheal cells) or GSGB9-1 (oenocytes) lines extends survival. We also found that the GSG3448 line had a lower tolerance to starvation (without RU486) (86.21% survival) after 24 hours than the GSGB9-1 (100% survival) lines (Fig. 5C,D).

**Figure 5.**
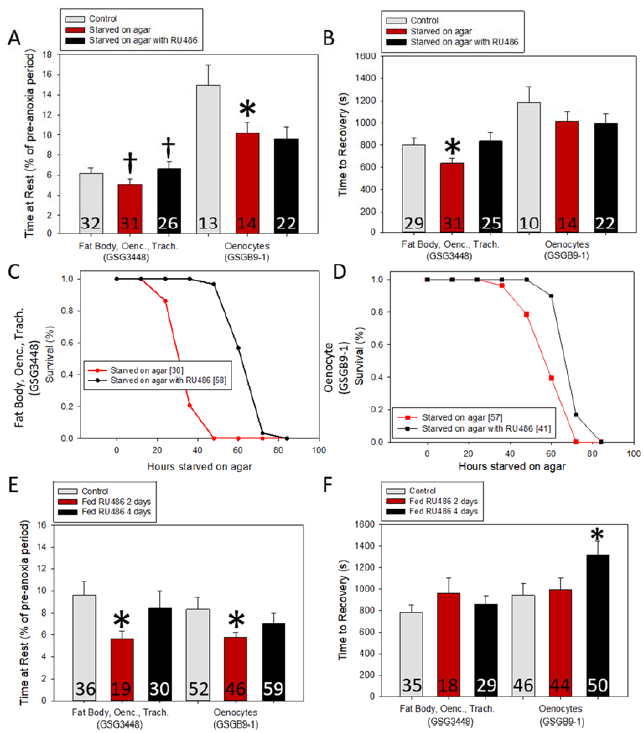
Effects of manipulation of AMPKα subunit expression in fat body and oenocytes with and without starvation. **(A,B)** Starvation (24 hr) induced hyperactivity in both the combined fat body, oenocyte, tracheal cell (GSG3448) and oenocyte (GSGB9-1) lines. Downregulation of the AMPKα subunit in the combined fat body, oenocyte, tracheal cell by adding 10 μg/mL RU486 to the agar increased TAR and blocked the effect of starvation on anoxic TR. p < 0.05 (*) by Mann-Whitney Rank Sum. **(C, D)** Survival curves of tissue-specific lines during starvation: agar alone (red line); agar with RU486 (black line). The combined fat body, oenocyte, tracheal cell line shows improved survival on agar with RU486 compared with the oenocyte line. **(E, F)** Both the combined fat body, oenocyte, tracheal cell and oenocyte lines had reduced TAR after 2 days fed RU486 but did not exhibit changes in anoxic TR. After four days fed RU486 both lines exhibit regular locomotor activity, however reduced AMPKα subunit affected the oenocyte line’s anoxic TR. p < 0.05 (*) by Mann-Whitney Rank Sum. Data are plotted as mean ± SE. Sample sizes are indicated in bars. Significance comparisons were made as (control) vs. (starvation on agar) or (starvation on agar) vs. (starvation on agar with RU486).

### AMPKα subunit affects starvation phenotype

If AMPKα was critical for one of the previously described starvation phenotypes - increased PL, reduced TAR, and fast TR - then its disruption using AMPKα-RNAi should abolish these effects. We first characterized each line's starvation phenotype. Starvation alone (without RU486) in AMPKα-RNAi Gene-Switch lines GSG3448 and GSGB9-1 caused a reduction in TAR (GSG3448: *Mann-Whitney, U* (30, 31) = 349, p = 0.095; GSGB9-1:*t-test, t* = 2.171, *df* = 24, p =0.020). Following anoxic exposure, the GSGB9-1 line showed a trend towards reducing TR (Table 1S), however only the combined expression line GSG3448 had a significantly reduced TR after 24 hrs of starvation (GSG3448: *Mann-Whitney, U* (29, 31) = 302.5, p = 0.030). This suggests, consistent with the results above, that these lines may have varying tolerance to starvation.

When feeding RU486 to inactivate AMPKα we found that only the combined expression line GSG3448 significantly affected the starvation phenotype. Starvation on agar with RU486 abolished the starvation-TAR phenotype; comparing starvation alone and starvation with RU486 (Fig. 5A) (*Mann-Whitney, U* (26, 31) = 296.5, p = 0.088). As well, starved GSG3448 flies no longer had a fast TR when fed RU486 (Fig. 5B) (*Mann-Whitney, U* (25, 31) = 291, p = 0.114). This suggests that during starvation the regulation of AMPKα through multiple tissues, either via a cooperative or an additive effect, may be required to induce a starvation phenotype.

However, as the GSG3448 line had a lower tolerance to starvation (Fig. 5C-E), we cannot rule out the possibility that AMPKα is simply upregulated more after 24 hrs of starvation than the single tissue line; resulting in a larger inhibitory effect of RU486.

### Short-term AMPKα subunit down-regulation affects locomotor behavior and tolerance to anoxia

To address the acute effects of reduced AMPK*α* subunit in the oenocyte and fat body, flies were fed RU486 for two and four days. We found that a two day downregulation of AMPK*α* in the GSG3448 and GSGB9-1 lines reduced TAR, which suggests a basal level of AMPK*α* may be active in these tissues and that the loss of AMPK*α* can modify locomotor behavior (Fig. 5E; GSG3448: *Mann-Whitney, U* (19, 36) = 181.5, p = 0.005; GSGB9-1: *Mann-Whitney, U* (46, 52) = 906, p = 0.039). However, flies fed RU486 for four days do not show this effect (GSG3448: *Mann-Whitney, U* (30, 36) = 430, p = 0.158; GSGB9-1: *Mann-Whitney, U* (52, 59) = 1290, p = 0.151). This return to regular locomotor behavior indicates that under normoxic conditions the fly may attempt to compensate for a loss of basal AMPK*α* within these tissues. However when faced with anoxic stress, AMPK*α* flies fed RU486 for four days took significantly longer to recover locomotor activity (TR: *Mann-Whitney, U* (46, 50) = 796, p = 0.010). This suggests mechanisms of basal AMPK*α* compensation were unable to help the fly during anoxic stress; which may require AMPK upregulation.

## Discussion

We investigated how food restriction affects the response of the nervous system to anoxia in adult *Drosophila*. Our main conclusions are: 1. Prior starvation speeds recovery of locomotion from an anoxic coma and this is mimicked by pharmacological treatments that would activate AMPK. 2. Collapse and restoration of ion gradients with anoxia/re-oxygenation are modulated by starvation and AMPK. This contributes little to the timing of recovery of locomotor circuit function, however, it could affect the susceptibility to SD. This remains to be determined. 3. Locomotor circuit recovery after anoxia is modulated by AMPK activity in glia, particularly the ensheathing glial subtype, rather than neurons. 4. AMPK in tissues that supply energy to the CNS (e.g. fat body, oenocytes and glia) can modulate locomotor circuit recovery and survival during periods of starvation.

We characterized a starvation phenotype of adult flies before, during, and after exposure to brief anoxia and contrasted this with the pharmacological activation of AMPK. We showed that when confined within a small arena, starving flies will modify their locomotor behavior to increase their distance travelled by spending less time inactive. A similar hyperactivity can be seen in flies fed metformin or AICAR. During anoxia, we found that starved flies are able to delay SD in the brain, however this does not affect the magnitude of SD or dynamics of the recovery of ion gradients. In contrast, flies fed metformin or AICAR display a smaller ionic disruption and prolonged recovery following SD. As both metformin and AICAR affect adenosine - (1) metformin decreases adenosine deaminase and (2) AICAR upon phosphorylation to ZMP shares structural similarities with adenosine - this finding parallels work in mammals which shows that SD impairs neurotransmission and that increased adenosine prolongs depression of electrocorticographic activity (Ouyang et al., 2011; Lindquist and Shuttleworth, 2017). Following anoxia, both pharmacological activation of AMPK and starvation improved locomotor recovery rates following anoxia. As both dietary restriction and pharmacological activators of AMPK have been previously shown to modulate oxidative stress in mammalian (Walsh et al., 2014; Shen et al., 2017) and non-mammalian organisms (Vigne et al., 2009; LaRue and Padilla, 2011), here we provide a tissue-specific evaluation of AMPK in modulating anoxic tolerance. We show that in the fly body, the oenocyte and fat body have independent and context-specific roles in anoxic tolerance, whereas glia is the sole AMPK-dependent tissue in the brain.

### Starvation, AMPK in the liver and oenocytes

Under fed conditions the fly fat body is involved in the steady transfer of lipids to the oenocytes (Brasaemle, 2007). In starved flies lipids accumulate in the oenocytes (Gutierrez et al., 2007). Lipid accumulation in the fly oenocytes is thought to function similar to a mammalian liver, whereby lipids are mobilized to a reserve pool to 'fuel' ketogenesis, a process which provides an alternative energy source to the brain during energetic stress. It has been suggested that ketone bodies act as a glucose-replacing fuel, during enhanced neuronal activity, as they are preferentially used for neuronal energy (Izumi et al., 1997; Blazquez et al., 1999). It has been previously reported that ketone bodies protect the brain from hypoxia and ischemic injury (Go et al., 1988; Auestad et al., 1991). Here we provide data that show, consistent with a pathway from lipid accumulation to neuroprotective ketogenesis, that starvation modifies anoxic tolerance.

As the 'metabolic load' of the mammalian liver is split between the oencotye and fat body in flies, we sought to determine whether AMPK in these tissues affects starvation-induced anoxic tolerance. Previous work investigating lipogenesis in *Drosophila* has shown that AMPK acts as a master switch for lipid regulation: AMPK upregulation, through the feeding of metformin, decreases total lipid content in the fly; and AMPK downregulation via the expression of a dominant negative alpha subunit variant, increases lipid content in the fly oencoytes (Johnson et al., 2010; Slack et al., 2012). Following this logic, decreasing AMPK during starvation should increase available lipid content for ketogenenic oxidation, and further increase anoxic tolerance. However, we found that when we disrupted AMPKα in the combined fat body, oenocytes, and tracheal cells line, but not the oenocyte line, the beneficial aspects of starvation are lost. This suggests that lipid accumulation is likely not the sole factor which allows starved individuals to tolerate anoxic bouts, but rather AMPK-dependent mechanisms in the fat body. AMPK has been shown to phosphorylate acetyl-CoA carboxylase (ACC1, ACC2) which turns off fatty acid synthesis and turns on fatty acid oxidation in an effort to conserve energy while producing ATP (Hardie, 2007). We therefore suggest that under global energetic stress, as induced by prolonged starvation, the AMPK-dependent shutdown of energetically expensive pathways outweighs the production of alternative catabolic pathways.

In unstressed, fed conditions the disruption of AMPKα in oenocytes had a time-dependent effect on tolerance to anoxic conditions. We report that downregulation of AMPKα for two days in either the combined fat body, oenocyte, and tracheal cell or oenocyte lines increased hyperactivity, and note that this trend was diminished by day four. Our observation is consistent with multiple observations that AMPK alters locomotor behavior (Lee and Park, 2004; Johnson et al., 2010; Ahmadi and Roy, 2016; Moller et al., 2016). We suspect that by day four regular behavior returns to normal due to the upregulation of other AMPK subunits, as has been seen in mice β2 KO mice (Steinberg et al., 2010), or through an independent compensatory pathway. Despite a return to regular locomotor behavior, we found that flies with reduced AMPKα in the oenocytes alone take a longer time to recover from anoxia; suggesting even in fed conditions a prolonged reduction of AMPKα in the oencotyes affects susceptibility to anoxic stresses.

### Neuronal AMPK in anoxic tolerance

While AMPK-driven ketogenesis is primarily thought to be derived from hepatic tissue, several studies have explored ketogenic machinery in neuronal tissue. In mammals AMPK is primarily expressed in neurons with some expression in astrocyte glia (Turnley et al., 1999; Culmsee et al., 2001). In particular, astrocyte glia has been shown to exhibit similar ketogenic machinery as hepatocytes (reviewed in (Guzman and Blazquez, 2004)): a preference for fatty acids over glucose as a primary metabolic fuel during ketogenesis (McGarry and Foster, 1980; Blazquez et al., 1998); carnitine palmitoyltransferase I (CPT-I) is the metabolic pace-setting step of ketogenesis (Drynan et al., 1996; Blazquez et al., 1998); similar ketogenic inhibitors, malonyl-CoA for CPT-I and acetyl-CoA carboxylase for malonyl-CoA, are observed in both astrocytes and hepatocytes (McGarry and Brown, 1997; Blazquez et al., 1998). However, as a primary signaling molecule, AMPK has also been shown to protect against neurotoxicity (Eom et al., 2016), increase neuronal survival following glucose deprivation or chemical hypoxia (Culmsee et al., 2001), mediate macrophage phagocytosis (Quan et al., 2015), and increase autophagy post-stroke to decrease infarct volume (Shen et al., 2017), amongst other findings. While we don't address the mechanisms which modulate anoxic tolerance in *Drosophila*, we do report, consistent with these findings, that upregulating AMPKα in glia affects locomotor recovery following anoxia. Further, to the best of our knowledge, our finding that AMPKα neural up- or down-regulation in *Drosophila* does not affect post-anoxia recovery has not been previously reported. Our data in combination with work in the fly showing glial but not neural Hsp70 mitigates loss of ion homeostatis during repetitive anoxic stress (Armstrong et al., 2011), suggest that in contrast to mammalian models, in *Drosophila,* glia is an effective target to manipulate anoxic tolerance in the fly brain.

In addition to evaluating neuronal AMPK up-regulation in anoxia tolerance in *Drosophila,* we suspected RNAi knockdown would prolong locomotor recovery times. First, we found that AMPK-RNAi was shown to be lethal in pan-glial but not pan-neural tissue during pupation. As AMPK inhibition was previously shown to block glycolysis in cultured mammalian astrocytes but not neurons (Funes et al., 2014), we were not surprised to find that others have shown dsRNA directed against components of the glycolytic pathway - trehalose, aldolase, and pyruvate - resulted in late larval and early pupal lethality in pan-glial but not pan-neural tissue in *Drosophila* (Volkenhoff et al., 2015). To avoid AMPK-RNAi pan-glial lethality, we targeted specific glial subtypes. Contrary to what we suspected, flies seemed to recover faster from anoxia. However, nearly all glial-specific knockdown lines had lower locomotory activity and higher body weight. This finding was again consistent with Volkenhoff *et al.'s* 2015 work, which showed that a knockdown of components in the glyolytic pathway that did not cause lethality showed severe locomotor defects when expressed in glia. Together these findings highlight the importance of the AMPK pathway in glia but not neurons for pupal development and locomotor behavior. Unfortunately, due to the likely changes in metabolism in these lines there is little we can say about AMPK disruption during anoxia without first uncoupling the existing phenotype changes. This work raises many questions as to not only what role glial AMPK plays in pupal development, but also how AMPK functions in these glial subtypes to regulate regular metabolic rate.

## Acknowledgements

We thank Laurent Seroude for his comments on the manuscript and Adam Chippindale for access to a Cahn-microbalance to weigh flies. Funded by a Discovery Grant to RMR from the Natural Sciences and Engineering Research Council of Canada.

## Supplementary Information

**Table 1S.** Summary of locomotor parameters for all tests. Due to the exclusion criteria following anoxia, the sample size for TR is smaller in certain cases. Statistical comparisons were made between their respective controls (as outlined in previous figures). p < 0.10 (ł), p < 0.05 (*), p < 0.01 (**), p < 0.001 (***). Details of statistical tests performed are included in the Results section.

|  | Treatment | n | Pre-anoxia Period |  | Anoxia | Post-anoxia Period |  |
| --- | --- | --- | --- | --- | --- | --- | --- |
|  |  |  | Path Length (cm) | Time at Rest (%) | Time to Succumb (s) | Time to Recovery (s) | n (TR) |
| Starvation | Starvation (control) | 32 | 215.3 ± 6.5 | 8.7 ± 2.1 | 10.5 ± 0.4 | 874.9 ± 82.0 | 25 |
|  | Starvation | 32 | 235.2 ± 5.3 * | 4.3 ± 0.4 * | 10.2 ± 0.4 | 631.6 ± 36.2 * | 32 |
| Pharmacology | Metformin (control) | 16 | 216.8 ± 6.5 | 12.0 ± 1.9 | 10.5 ± 0.4 | 1178.8 ± 151.3 | 16 |
|  | Metformin | 16 | 252.0 ± 8.2 ** | 7.3 ± 0.8 * | 10.3 ± 0.5 | 823.2 ± 69.8 † | 16 |
|  | AICAR (control) | 32 | 221.3 ± 4.0 | 7.7 ± 0.8 | 8.4 ± 0.1 | 855.7 ± 49.2 | 32 |
|  | AICAR | 32 | 235.5 ± 5.5 * | 6.7 ± 0.6 | 8.3 ± 0.1 | 710.2 ± 43.3 * | 32 |
| Driver | Genotype |  |  |  |  |  |  |
| Pan-neural | 8765/+ | 24 | 219.4 ± 10.6 | 10.1 ± 2.4 | 8.0 ± 0.1 | 864.8 ± 88.2 | 24 |
|  | 8765/ UAS-Cherry-AMPKα | 24 | 208.8 ± 7.2 | 6.5 ± 0.7 | 8.3 ± 0.2 | 1030.7 ± 146.0 | 24 |
|  | 8765/UAS-AMPK-RNAi | 15 | 203.3 ± 8.8 | 6.1 ± 1.5 | 8.3 ± 0.3 | 717.2 ± 55.9 | 15 |
| Pan-glial | 7415/+ | 24 | 240.5 ± 8.0 | 7.90 ± 1.2 | 8.1 ± 0.1 | 770.1 ± 108.7 | 24 |
|  | 7415/UAS-Cherry-AMPKα | 24 | 227.7 ± 9.4 | 8.5 ± 0.9 | 7.8 ± 0.09 | 634.3 ± 77.6 * | 24 |
| Upstream Sequence | UAS-Cherry-AMPKα/+ | 24 | 250.1 ± 7.2 | 4.6 ± 0.5 | 8.8 ± 0.4 | 703.1 ± 43.7 | 24 |
|  | UAS-AMPKα-RNAi/+ | 23 | 274.3 ± 7.8 | 5.9 ± 1.3 | 8.2 ± 0.1 | 994.9 ± 78.2 | 23 |
| Glial Class Driver | Genotype |  |  |  |  |  |  |
| Subperineural | NP2276/+ | 32 | 255.7 ± 12.7 | 5.4 ± 0.5 | 9.1 ± 0.1 | 906.5 ± 104.1 | 32 |
|  | NP2276/UAS-AMPKα-RNAi | 28 | 183.6 ± 13.5 * | 8.1 ± 2.0 | 8.6 ± 0.5 | 837.1 ± 88.6 | 26 |
| Astrocyte | NP1243/+ | 48 | 158.2 ± 9.2 | 13.3 ± 1.3 | 8.5 ± 0.4 | 929.3 ± 62.1 | 47 |
|  | NP1243/UAS-AMPKα-RNAi | 47 | 151.3 ± 9.8 | 8.5 ± 0.6 | 8.3 ± 0.3 | 813.5 ± 72.6 | 45 |
| Enstheating | NP6520/+ | 64 | 214.3 ± 10.7 | 6.1 ± 0.5 | 8.5 ± 0.2 | 548.8 ± 25.6 | 64 |
|  | NP6520/UAS-AMPKα-RNAi | 63 | 164.7 ± 8.6 *** | 6.2 ± 0.6 | 7.6 ± 0.3 * | 419.0 ± 19.7 * | 63 |
|  | MZ0709/+ | 64 | 204.7 ± 10.4 | 9.3 ± 0.8 | 8.9 ± 0.2 | 683.0 ± 40.7 | 64 |
|  | MZ0709/UAS-AMPKα-RNAi | 66 | 157.6 ± 7.3 * | 5.8 ± 0.5 | 8.2 ± 0.3 | 503.2 ± 13.2 | 66 |
|  | Alrm II/+ | 56 | 218.6 ± 12.8 | 4.0 ± 0.4 | 9.2 ± 0.2 | 744.0 ± 47.3 | 56 |
|  | Alrm II/UAS-AMPKα-RNAi | 51 | 175.7 ± 10.2 *** | 5.1 ± 0.6 | 8.0 ± 0.3 | 808.2 ± 83.8 | 47 |
|  | Alrm III/+ | 63 | 191.9 ± 10.9 | 7.0 ± 0.9 | 9.0 ± 0.1 | 849.6 ± 57.3 | 58 |
|  | Alrm III/UAS-AMPKα-RNAi | 48 | 152.2 ± 13.2 *** | 11.5 ± 1.7 * | 8.2 ± 0.3 | 641.8 ± 49.5 | 48 |
| Upstream Sequence | UAS-AMPKα-RNAi/+ | 111 | 201.4 ± 8.1 | 5.5 ± 0.5 | 8.6 ± 0.2 | 695.3 ± 49.7 | 109 |
| GeneSwitch Lines | Genotype (Driver/UAS-AMPKα-RNAi) & Treatment |  |  |  |  |  |  |
| Fat body, oenocytes, and tracheal cells | GSG3448 (control) | 32 | 174.2 ± 4.4 | 6.2 ± 0.5 | 7.5 ± 0.2 | 796.9 ± 71.4 | 29 |
|  | GSG3448 (starved on agar) | 31 | 173.8 ± 4.1 | 5.1 ± 0.6 † | 7.7 ± 0.2 | 639.2 ± 43.6 * | 31 |
|  | GSG3448 (starved on agar with RU486) | 26 | 175.5 ± 5.8 | 6.6 ± 0.7 † | 7.8 ± 0.2 | 836.3 ± 80.2 † | 25 |
| Oenocytes | GSGB9-1 (control) | 13 | 121.5 ± 8.9 | 15.0 ± 2.0 | 6.3 ± 0.8 | 1180.1 ± 144.6 | 10 |
|  | GSGB9-1 (starved on agar) | 14 | 118.2 ± 5.8 * | 10.2 ± 1.1 * | 6.6 ± 0.9 | 1017.9 ± 85.1 | 14 |
|  | GSGB9-1 (starved on agar with RU486) | 22 | 131.0 ± 5.2 † | 9.6 ± 1.2 | 7.6 ± 0.4 | 994.9 ± 88.9 | 22 |
| Fat body | GSG10751 (control) | 14 | 144.7 ± 7.4 | 11.2 ± 3.5 | 8.6 ± 0.2 | 1153.5 ± 158.8 | 13 |
|  | GSG10751 (starved on agar) | 20 | 124.3 ± 5.2 | 14.3 ± 2.0 * | 7.5 ± 0.6 | 933.0 ± 82.4 | 20 |
|  | GSG10751 (starved on agar with RU486) | 16 | 109.3 ± 6.4 † | 20.2 ± 3.7 | 6.5 ± 0.5 * | 1102.5 ± 155.3 | 15 |
| Fat body, oenocytes, and tracheal cells | GSG3448 (control) | 36 | 160.6 ± 3.7 | 8.3 ± 0.7 | 7.2 ± 0.3 | 784.5 ± 68.7 | 35 |
|  | GSG3448 (fed RU486 2 days) | 19 | 165.2 ± 5.7 | 5.6 ± 0.7 | 8.0 ± 0.4 | 960.4 ± 139.6 | 18 |
|  | GSG3448 (fed RU486 4 days) | 30 | 153.6 ± 5.3 † | 8.4 ± 1.6 | 6.7 ± 0.3 † | 857.0 ± 78.8 | 29 |
| Oenocytes | GSGB9-1 (control) | 52 | 159.6 ± 3.8 | 8.3 ± 1.1 | 7.4 ± 0.2 | 942.0 ± 106.2 | 46 |
|  | GSGB9-1 (fed RU486 2 days) | 46 | 160.7 ± 3.5 | 5.8 ± 0.5 * | 7.5 ± 0.1 | 993.9 ± 106.0 | 44 |
|  | GSGB9-1 (fed RU486 4 days) | 59 | 161.7 ± 4.3 | 7.1 ± 0.9 | 7.4 ± 0.2 | 1319.7 ± 124.7 † | 50 |
| Fat body | GSG10751 (control) | 38 | 165.6 ± 4.1 | 6.6 ± 0.5 | 7.8 ± 0.2 | 736.2 ± 64.3 | 36 |
|  | GSG10751 (fed RU486 2 days) | 30 | 183.2 ± 6.9 † | 5.4 ± 0.5 | 7.8 ± 0.4 | 605.1 ± 57.5 † | 30 |
|  | GSG10751 (fed RU486 4 days) | 40 | 171.8 ± 4.3 † | 6.4 ± 0.6 † | 8.0 ± 0.1 | 783.2 ± 63.5 | 39 |

**Video 1S. Locomotor activity before, during and after anoxia in fed and starved adult flies.** The video shows four columns of eight locomotor assay chambers (1.27 cm diameter) each containing a single fly. The first two columns contain flies that have been fed (Control); the second two columns contain flies that have been maintained without food, but with water, for 24 hours (Starved). The first 30 s of the video shows 5 minutes of activity with playback at 10x the acquisition rate. The next 30 s is played at normal speed for the duration of the nitrogen application. Note the animals entering a coma after 10 – 15 s of anoxia. The final 3 mins shows 30 minutes of playback at 10x speed to show post-anoxic recovery of locomotor function. Note that starved flies are more active during the baseline activity and recover from anoxia faster than control flies.

## References

Ahmadi M, Roy R (2016) AMPK acts as a molecular trigger to coordinate glutamatergic signals and adaptive behaviours during acute starvation. Elife 5:e16349.

Armstrong GA, Xiao C, Krill JL, Seroude L, Dawson-Scully K, Robertson RM (2011) Glial Hsp70 protects K+ homeostasis in the *Drosophila* brain during repetitive anoxic depolarization. PLoS One 6:e28994.

Auestad N, Korsak RA, Morrow JW, Edmond J (1991) Fatty acid oxidation and ketogenesis by astrocytes in primary culture. J Neurochem 56:1376–1386.

Awasaki T, Lai SL, Ito K, Lee T (2008) Organization and postembryonic development of glial cells in the adult central brain of *Drosophila*. J Neurosci 28:13742–13753.

Benasayag-Meszaros R, Risley MG, Hernandez P, Fendrich M, Dawson-Scully K (2015) Pushing the limit: Examining factors that affect anoxia tolerance in a single genotype of adult *D. melanogaster*. Scientific reports 5:9204.

Bland ML, Lee RJ, Magallanes JM, Foskett JK, Birnbaum MJ (2010) AMPK supports growth in *Drosophila* by regulating muscle activity and nutrient uptake in the gut. Dev Biol 344:293–303.

Blazquez C, Sanchez C, Velasco G, Guzman M (1998) Role of carnitine palmitoyltransferase I in the control of ketogenesis in primary cultures of rat astrocytes. J Neurochem 71:1597– 1606.

Blazquez C, Woods A, de Ceballos ML, Carling D, Guzman M (1999) The AMP-activated protein kinase is involved in the regulation of ketone body production by astrocytes. J Neurochem 73:1674–1682.

Brasaemle DL (2007) Thematic review series: adipocyte biology. The perilipin family of structural lipid droplet proteins: stabilization of lipid droplets and control of lipolysis. J Lipid Res 48:2547–2559.

Callier V, Hand SC, Campbell JB, Biddulph T, Harrison JF (2015) Developmental changes in hypoxic exposure and responses to anoxia in *Drosophila melanogaster*. J Exp Biol 218:2927–2934.

Culmsee C, Monnig J, Kemp BE, Mattson MP (2001) AMP-activated protein kinase is highly expressed in neurons in the developing rat brain and promotes neuronal survival following glucose deprivation. J Mol Neurosci 17:45–58.

de Crespigny AJ, Rother J, Beaulieu C, Moseley ME, Hoehn M (1999) Rapid monitoring of diffusion, DC potential, and blood oxygenation changes during global ischemia. Effects of hypoglycemia, hyperglycemia, and TTX. Stroke 30:2212–2222.

Dreier JP, Reiffurth C (2015) The stroke-migraine depolarization continuum. Neuron 86:902– 922.

Drynan L, Quant PA, Zammit VA (1996) Flux control exerted by mitochondrial outer membrane carnitine palmitoyltransferase over beta-oxidation, ketogenesis and tricarboxylic acid cycle activity in hepatocytes isolated from rats in different metabolic states. Biochem J 317:791–795.

Eom JW, Lee JM, Koh JY, Kim YH (2016) AMP-activated protein kinase contributes to zinc-induced neuronal death via activation by LKB1 and induction of Bim in mouse cortical cultures. Mol Brain 9:14.

Funes HA, Apostolova N, Alegre F, Blas-Garcia A, Alvarez A, Marti-Cabrera M, Esplugues JV (2014) Neuronal bioenergetics and acute mitochondrial dysfunction: a clue to understanding the central nervous system side effects of efavirenz. J Infect Dis 210:1385– 1395.

Galli GL, Richards JG (2014) Mitochondria from anoxia-tolerant animals reveal common strategies to survive without oxygen. J Comp Physiol B 184:285–302.

Gibson CL, Murphy AN, Murphy SP (2012) Stroke outcome in the ketogenic state--a systematic review of the animal data. J Neurochem 123 Suppl 2:52–57.

Go KG, Prenen GH, Korf J (1988) Protective effect of fasting upon cerebral hypoxic-ischemic injury. Metab Brain Dis 3:257–263.

Gutierrez E, Wiggins D, Fielding B, Gould AP (2007) Specialized hepatocyte-like cells regulate *Drosophila* lipid metabolism. Nature 445:275–280.

Guzman M, Blazquez C (2004) Ketone body synthesis in the brain: possible neuroprotective effects. Prostaglandins Leukot Essent Fatty Acids 70:287–292.

Hansen AJ (1985) Effect of anoxia on ion distribution in the brain. Physiol Rev 65:101–148.

Hardie DG (2007) AMP-activated/SNF1 protein kinases: conserved guardians of cellular energy. Nat Rev Mol Cell Biol 8:774–785.

Hardie DG, Carling D, Carlson M (1998) The AMP-activated/SNF1 protein kinase subfamily: metabolic sensors of the eukaryotic cell? Annu Rev Biochem 67:821–855.

Hardie DG, Salt IP, Hawley SA, Davies SP (1999) AMP-activated protein kinase: an ultrasensitive system for monitoring cellular energy charge. Biochem J 338 (Pt 3):717-722.

Hawley SA, Selbert MA, Goldstein EG, Edelman AM, Carling D, Hardie DG (1995) 5'-AMP activates the AMP-activated protein kinase cascade, and Ca2+/calmodulin activates the calmodulin-dependent protein kinase I cascade, via three independent mechanisms. J Biol Chem 270:27186–27191.

Hochachka PW, Buck LT, Doll CJ, Land SC (1996) Unifying theory of hypoxia tolerance: molecular/metabolic defense and rescue mechanisms for surviving oxygen lack. Proc Natl Acad Sci U S A 93:9493–9498.

Izumi Y, Benz AM, Katsuki H, Zorumski CF (1997) Endogenous monocarboxylates sustain hippocampal synaptic function and morphological integrity during energy deprivation. J Neurosci 17:9448–9457.

Jibb LA, Richards JG (2008) AMP-activated protein kinase activity during metabolic rate depression in the hypoxic goldfish, *Carassius auratus*. J Exp Biol 211:3111–3122.

Johnson EC, Kazgan N, Bretz CA, Forsberg LJ, Hector CE, Worthen RJ, Onyenwoke R, Brenman JE (2010) Altered metabolism and persistent starvation behaviors caused by reduced AMPK function in *Drosophila*. PLoS One 5:e12799.

LaRue BL, Padilla PA (2011) Environmental and genetic preconditioning for long-term anoxia responses requires AMPK in *Caenorhabditis elegans*. PLoS One 6:e16790.

Lee G, Park JH (2004) Hemolymph sugar homeostasis and starvation-induced hyperactivity affected by genetic manipulations of the adipokinetic hormone-encoding gene in *Drosophila melanogaster*. Genetics 167:311–323.

Lee JH, Koh H, Kim M, Kim Y, Lee SY, Karess RE, Lee SH, Shong M, Kim JM, Kim J, Chung J (2007) Energy-dependent regulation of cell structure by AMP-activated protein kinase. Nature 447:1017–1020.

Li J et al. (2016) An obligatory role for neurotensin in high-fat-diet-induced obesity. Nature 533:411–415.

Lighton JR, Schilman PE (2007) Oxygen reperfusion damage in an insect. PLoS One 2:e1267.

Lindquist BE, Shuttleworth CW (2017) Evidence that adenosine contributes to Leao's spreading depression in vivo. J Cereb Blood Flow Metab 37:1656–1669.

Mantovani J, Roy R (2011) Re-evaluating the general(ized) roles of AMPK in cellular metabolism. FEBS Lett 585:967–972.

Manwani B, McCullough LD (2013) Function of the master energy regulator adenosine monophosphate-activated protein kinase in stroke. J Neurosci Res 91:1018–1029.

McGarry JD, Foster DW (1980) Regulation of hepatic fatty acid oxidation and ketone body production. Annu Rev Biochem 49:395–420.

McGarry JD, Brown NF (1997) The mitochondrial carnitine palmitoyltransferase system. From concept to molecular analysis. Eur J Biochem 244:1–14.

Moller LL, Sylow L, Gotzsche CR, Serup AK, Christiansen SH, Weikop P, Kiens B, Woldbye DP, Richter EA (2016) Decreased spontaneous activity in AMPK alpha2 muscle specific kinase dead mice is not caused by changes in brain dopamine metabolism. Physiol Behav 164:300–305.

Money TGA, Sproule MKJ, Hamour AF, Robertson RM (2014) Reduction in neural performance following recovery from anoxic stress Is mimicked by AMPK pathway activation. PLoS One 9:e88570.

Murray AJ, Knight NS, Cole MA, Cochlin LE, Carter E, Tchabanenko K, Pichulik T, Gulston MK, Atherton HJ, Schroeder MA, Deacon RM, Kashiwaya Y, King MT, Pawlosky R, Rawlins JN, Tyler DJ, Griffin JL, Robertson J, Veech RL, Clarke K (2016) Novel ketone diet enhances physical and cognitive performance. Faseb J 30:4021–4032.

Osterwalder T, Yoon KS, White BH, Keshishian H (2001) A conditional tissue-specific transgene expression system using inducible GAL4. Proc Natl Acad Sci U S A 98:12596– 12601.

Ouyang J, Parakhia RA, Ochs RS (2011) Metformin activates AMP kinase through inhibition of AMP deaminase. J Biol Chem 286:1–11.

Quan H, Kim JM, Lee HJ, Lee SH, Choi JI, Bae HB (2015) AICAR Enhances the phagocytic ability of macrophages towards apoptotic cells through P38 mitogen activated protein kinase activation independent of AMP-Activated protein kinase. PLoS One 10:e0127885.

Rodgers-Garlick CI, Armstrong GAB, Robertson RM (2011) Metabolic stress modulates motor patterning via AMP-activated protein kinase. J Neurosci 31:3207–3216.

Rodriguez EC, Robertson RM (2012) Protective effect of hypothermia on brain potassium homeostasis during repetitive anoxia in *Drosophila melanogaster*. Journal of Experimental Biology 215:4157–4165.

Schulz JG, Laranjeira A, Van Huffel L, Gartner A, Vilain S, Bastianen J, Van Veldhoven PP, Dotti CG (2015) Glial beta-oxidation regulates *Drosophila* energy metabolism. Sci Rep 5:7805.

Shen P, Hou S, Zhu M, Zhao M, Ouyang Y, Feng J (2017) Cortical spreading depression preconditioning mediates neuroprotection against ischemic stroke by inducing AMP-activated protein kinase-dependent autophagy in a rat cerebral ischemic/reperfusion injury model. J Neurochem 140:799–813.

Slack C, Foley A, Partridge L (2012) Activation of AMPK by the putative dietary restriction mimetic metformin is insufficient to extend lifespan in *Drosophila*. PLoS One 7:e47699.

Spong KE, Andrew RD, Robertson RM (2016a) Mechanisms of spreading depolarization in vertebrate and insect central nervous systems. J Neurophysiol 116:1117–1127.

Spong KE, Rodriguez EC, Robertson RM (2016b) Spreading depolarization in the brain of *Drosophila* is induced by inhibition of the Na+/K+-ATPase and mitigated by a decrease in activity of protein kinase G. J Neurophysiol 116:1152–1160.

Staples JF, Buck LT (2009) Matching cellular metabolic supply and demand in energy-stressed animals. Comp Biochem Physiol A Mol Integr Physiol 153:95–105.

Steinberg GR, O'Neill HM, Dzamko NL, Galic S, Naim T, Koopman R, Jorgensen SB, Honeyman J, Hewitt K, Chen ZP, Schertzer JD, Scott JW, Koentgen F, Lynch GS, Watt MJ, van Denderen BJ, Campbell DJ, Kemp BE (2010) Whole body deletion of AMP-activated protein kinase {beta}2 reduces muscle AMPK activity and exercise capacity. J Biol Chem 285:37198–37209.

Suzuki M, Suzuki M, Sato K, Dohi S, Sato T, Matsuura A, Hiraide A (2001) Effect of beta-hydroxybutyrate, a cerebral function improving agent, on cerebral hypoxia, anoxia and ischemia in mice and rats. Jpn J Pharmacol 87:143–150.

Swick LL, Kazgan N, Onyenwoke RU, Brenman JE (2013) Isolation of AMP-activated protein kinase (AMPK) alleles required for neuronal maintenance in *Drosophila melanogaster*. Biol Open 2:1321–1323.

Turnley AM, Stapleton D, Mann RJ, Witters LA, Kemp BE, Bartlett PF (1999) Cellular distribution and developmental expression of AMP-activated protein kinase isoforms in mouse central nervous system. J Neurochem 72:1707–1716.

Ulgherait M, Rana A, Rera M, Graniel J, Walker DW (2014) AMPK modulates tissue and organismal aging in a non-cell-autonomous manner. Cell Rep 8:1767–1780.

Vigne P, Tauc M, Frelin C (2009) Strong dietary restrictions protect *Drosophila* against anoxia/reoxygenation injuries. PLoS One 4:e5422.

Volkenhoff A, Weiler A, Letzel M, Stehling M, Klambt C, Schirmeier S (2015) Glial glycolysis is essential for neuronal survival in *Drosophila*. Cell Metab 22:437–447.

Walsh ME, Shi Y, Van Remmen H (2014) The effects of dietary restriction on oxidative stress in rodents. Free Radic Biol Med 66:88–99.

Xiao C, Robertson RM (2015) Locomotion induced by spatial restriction in adult *Drosophila*. PLoS One 10:e0135825

Xiao C, Robertson RM (2016) Timing of locomotor recovery from anoxia modulated by the *white* gene in *Drosophila*. Genetics 203:787–797.

Zhu CD, Wang ZH, Yan B (2013) Strategies for hypoxia adaptation in fish species: a review. J Comp Physiol B 183:1005–1013.

